# Selfing-aware demographic inference highlights local population structure and bottlenecks in Hawaiian *Caenorhabditis elegans*

**DOI:** 10.64898/2026.09.22.753446

**Authors:** Chenxi Wang, Robyn E. Tanny, Konrad Lohse, Erik C. Andersen, Matthew Hartfield

## Abstract

Demographic inference based on sequentially Markovian coalescent (SMC) model is widely used for disentangling the evolutionary history of species and populations. However, the genetic effects of self-fertilization, including the expected reduction of population-level genetic diversity and hence effective population size (*N*_*e*_), are often ignored in demographic inference, despite the fact that selfing is a common reproductive mode in both invertebrates and plants. To investigate how self-fertilization could bias demographic inference, we apply eSMC2, a selfing-aware demographic inference method, to Hawaiian populations of *Caenorhabditis elegans*, a predominantly self-fertilizing nematode. Using an expanded dataset that contains more Hawaiian strains than in previous studies, we recovered previously reported genetic groups, including one largely consisting of strains from the Big Island. Population substructure is also detected among these Big Island subpopulations, highlighting the importance of identifying subgroups to avoid misinterpretations of demography. For different Big Island subpopulations, we estimate high selfing rates compatible with previous studies and infer consistent population declines. Population sizes inferred using eSMC2 are smaller than those inferred using PSMC’, which does not account for selfing. However, rescaling PSMC’ parameters to account for self-fertilization produces a similar demographic trajectory as eSMC2 inference. Our study thus demonstrates how one can account for effects of selfing when using SMC-based methods. It also indicates that the Hawaiian group shows population declines similar to those previously found outside Hawaii.

## Introduction

Information from sampled genomic data can be used to infer historical processes, including population size changes, migration patterns and divergence times (Marchi et al. 2021). Inferring the past demographic history of populations informs us of its evolutionary history, such as expansions, contractions, bottlenecks and colonization of new habitats (Sellinger et al. 2020), and also provides a baseline model for inferring selection (Johri et al. 2020, 2021; Soni and Jensen 2025). Coalescent theory underpins most demographic inference methods, which models the genealogical relationships of sampled sequences backwards in time (Kingman 1982). Here, the coalescence probability per unit of time (or coalescent rate) between a pair of sequences is inversely proportional to the population size in an ideal population under several assumptions, including random mating, no migration, no population structure, and no selection (Wakeley 2008). In real populations, the *coalescent effective population size* (*N*_*e*_) is the value that is actually inferred from genetic data (Sjödin et al. 2005). It refers to the size of an idealized population with the same probability of coalescence (i.e., the same level of genetic drift) as the actual population (Wright 1931; Crow 2010). Although coalescent *N*_*e*_ is an important indicator of past demography, the coalescent rate inferred under the population genetics framework is not only affected by population sizes but also other factors such as selection and population structure (Mazet et al. 2016; Boitard et al. 2022).

The coalescent rate is also affected by life-history traits of the species, including a population’s mating systems (Sellinger *et al*. 2020; Strütt *et al*. 2023; Peede *et al*. 2025). Self-fertilization (or selfing) is one of the most common non-random mating systems, in which two gametes coming from the same individual are involved in fertilization. About 5–6% of animal species are hermaphroditic (increasing to one third if excluding insects) and more than 60% of them exhibit an appreciable level (outcrossing rate less than 0.8) of selfing when assuming selfing is the only source of inbreeding (Jarne and Auld 2006). In plants, around 50% of species have an outcrossing rate less than 0.8 indicated by similar estimates (Goodwillie *et al*. 2005). The genetic consequences of selfing includes reduced individual heterozygosity and population-level polymorphism. Therefore, selfing increases the coalescent rate of alleles within the same individual, lowering *N*_*e*_ (Nordborg and Donnelly 1997; Cutter 2019). It also reduces the effective recombination rate, which causes longer spans of linkage disequilibrium along the genome and thus extends the effect of linked selection (Golding and Strobeck 1980; Nordborg 2000; Cutter 2019). Ignoring these genetic effects is likely to cause a misinterpretation of the inferred *N*_*e*_ changes.

The Sequentially Markovian coalescent (SMC) model underlies several methods for inferring demographic history, which models the observed genotypes along the sequence as a simple Markovian structure for inferring population genetic parameters (McVean and Cardin 2005). A set of popular demographic inference methods have been derived from the model, including PSMC (Li and Durbin 2011), MSMC (Schiffels and Durbin 2014), and SMC++ (Terhorst et al. 2017). However, most SMC-based methods do not account for the effect of self-fertilization. One exception is eSMC (Sellinger et al. 2020), which was developed for a joint inference of both a constant selfing rate (the proportion of offspring produced through selfing) and demographic history. eSMC adopts the framework of PSMC’ (i.e., MSMC using only two haplotypes) with rescaled parameters to account for selfing or seed-banking (Sellinger et al. 2020). Based on the coalescent with partial selfing model described in Nordborg and Donnelly (1997) and Nordborg (2000), it considers how coalescent and recombination rates are rescaled differently by selfing to obtain less-biased estimates of population sizes (Sellinger et al. 2020). It also adopts likelihood optimisation under constraints by allowing the user to set boundaries to parameters to be inferred, which can potentially reduce the variability of inference. Its accuracy under a wide range of demographic and selfing rate scenarios was validated by simulated data. Applying the method to *Arabidopsis thaliana*, a highly selfing plant (with selfing rate estimates ranging from around 90% to over 99%; Redei (1975); Abbott and Gomes (1989); Bakker *et al*. (2006); Picó *et al*. (2008); Platt *et al*. (2010)), infers a selfing rate of *∼* 87% and recent population declines in Swedish and German samples (Sellinger et al. 2020).

*Caenorhabditis elegans* is an androdioecious species with most individuals being hermaphrodites. Males occur in an extremely low rate in the wild (Barrière and Félix 2005a) and both recombination-based and heterozygote-based measures support a very low outcrossing rate, close to or even lower than 1% (Barrière and Félix 2005a; Cutter 2006; Barrière and Félix 2007). Strains isolated from the same location can exhibit high genetic identity due to the selfing nature of the species. Therefore, wild strains that are nearly identical are usually classified to isotypes (i.e., distinct genomewide genotypes) (Andersen et al. 2012; Lee et al. 2021; Crombie et al. 2023b). Previous population-level sequencing studies reveal a higher level of genetic diversity in the Hawaiian population compared to other regions of the globe, which are dominated by a set of commonly shared haplotypes possibly generated by chromosome-scale selective sweeps (Andersen et al. 2012; Crombie et al. 2019; Lee et al. 2021). The ‘Out of Hawaii’ hypothesis proposes that populations outside Hawaii originated from recent long-range dispersal or transfer events from a Hawaiian source population (Crombie et al. 2019). To validate this hypothesis, it is essential to investigate whether the Hawaiian population shared the same demographic history as other populations. However, previous *C. elegans* demography history studies were hampered by limited samples, especially from the Hawaiian Islands. For example, the *C. elegans* dataset used in Thomas *et al*. (2015) contains only one Hawaiian strain, which only allowed them to infer the split time between Hawaiian and other populations, rather than the demographic history of the Hawaiian population itself. Teterina *et al*. (2023) inferred a precipitous decline in *N*_*e*_ for samples from the Hawaiian Islands using SMC++, but all samples used in the study are from the ‘global’ genetic cluster identified in Lee *et al*. (2021). As the ‘global’ cluster consists largely of isotypes from outside the Hawaiian Islands, the past demography inferred in Teterina *et al*. (2023) may reflect the demographic history of strains that are not Hawaiian natives. As Hawaiian populations are less affected by genome-wide selective sweeps (Andersen et al. 2012; Crombie *et al*. 2019; Lee et al. 2021), their inferred demographic history could also be least biased by extensive linked selection. Finally, the possible ancient origin of selfing allows us to infer a demograhic history assuming a constant selfing rate. The estimated divergence time between *C. elegans* and its outcrossing sister-species *C. inopinata* is more than 100 million generations (Kanzaki *et al*. 2018). Previous population genetics and molecular evolution studies have provided several loose time estimates of the age of selfing (Cutter 2008; Cutter *et al*. 2008; Rane *et al*. 2010), with the lowest one being 23.3 million generations ago (Cutter *et al*. 2008).

In this study, we used a large whole-genome sequencing dataset of *C. elegans* to determine population structure and demographic history in this species, with a focus on Hawaiian populations. We study strains from CaeNDR (Crombie *et al*. 2023b) that contains more Hawaiian samples than used in previously studies, to first identify population structure and genetically distinct populations. We then focus on inferring the selfing rate and past demography of several populations constituting the ‘Hawaiian’ group reported in Lee *et al*. (2021) using eSMC2, the most recent version of eSMC. We also compared eSMC2 inference with PSMC’ to examine how eSMC2 accounts for selfing, and if non selfing-aware methods can be adapted for use with selfing species.

## Materials and Methods

### Dataset description

Variant call format (VCF) data for 611 *C. elegans* isotype reference strains representing 611 isotypes were downloaded from CaeNDR (Crombie *et al*. 2023b) (20231213 release, Crombie *et al*. (2023a)), where the reference *C. elegans* genome (WS283 release of Wormbase) and detailed information of all isotype reference strains are available. Compared to Lee *et al*. (2021) which identified 45 isotypes from the Hawaiian Islands, 203 Hawaiian isotype reference strains are included in this release, with 55 strains from Kauai, six strains from Oahu, two strains from Molokai, 29 strains from Maui and 111 strains from the Big Island. As *C. elegans* is a predominantly self-fertilizing species, heterozygous single nucleotide variant (SNV) calls are most likely errors. As documented in the CaeNDR Methods (Crombie *et al*. 2023a), ‘heterozygous SNV polarization’ was applied to the initial genotype calls to eliminate genotype calling errors that create false heterozygous sites. Here, polarization means converting biallelic heterozygous SNVs with sufficient evidence for conversion to homozygous genotypes. Then, the dataset was soft-filtered by flagging low-quality sites at both the site level and the sample level but without modifying or removing them. For the hard-filtered dataset, low-quality sites at the site level were removed and low-quality sites at the sample level were converted to missing.

### Heterozygous site check

To investigate whether variants are homozygous enough to allow considering strains as haploids, we calculated the count of heterozygous strains per site both before and after polarization, and with or without the site-level hard filter used by CaeNDR (except for the heterozygous fraction filter, as our aim was to compare the heterozygous rate of different datasets; and the missingness filter, which is discussed below). Specifically, we reverted polarized homozygote SNVs in the soft-filtered dataset back to heterozygous calls using the setGT plugin of BCFtools v.1.19 (Li 2011) and extracted genotypes using `bcftools query`. These genotypes were used to calculate the count of heterozygous strains at each site using a customized script (available on the accompanying Github repository).

There are 6,620,075 variants in total in this release of the soft-filtered dataset (Supplementary Table 1). Before heterozygous SNV polarization, 4,113,931 (62.14%) variants have one or more heterozygous genotypes but most of them are called heterozygous in fewer than 10 strains (Supplementary Fig. 1). After polarization, 1,317,649 (19.90% of the total) variants have at least one heterozygous genotype converted to homozygous genotype and the proportion of sites with one or more heterozygous genotypes decreases to 56.44%. 7,105,545 heterozygous calls are converted to homozygous calls and the mean number of converted genotypes per site is 1.07 out of 611 strains, which means the mean heterozygous strain count per site drops from 7.47 to 6.40 (Supplementary Table 1).

**Table 1.** Weighted average recombination rates of each chromosome and all autosomes in Caenorhabditis elegans based on (Rockman and Kruglyak 2009).

| Chromosome | Weighted average recombination rate<br>(crossover/bp/generation) |
| --- | --- |
| I | $3.12 \times 10^{-8}$ |
| II | $3.53 \times 10^{-8}$ |
| III | $3.91 \times 10^{-8}$ |
| IV | $2.71 \times 10^{-8}$ |
| V | $2.47 \times 10^{-8}$ |
| X | $2.95 \times 10^{-8}$ |
| Autosomes | $3.08 \times 10^{-8}$ |

**Figure 1.**
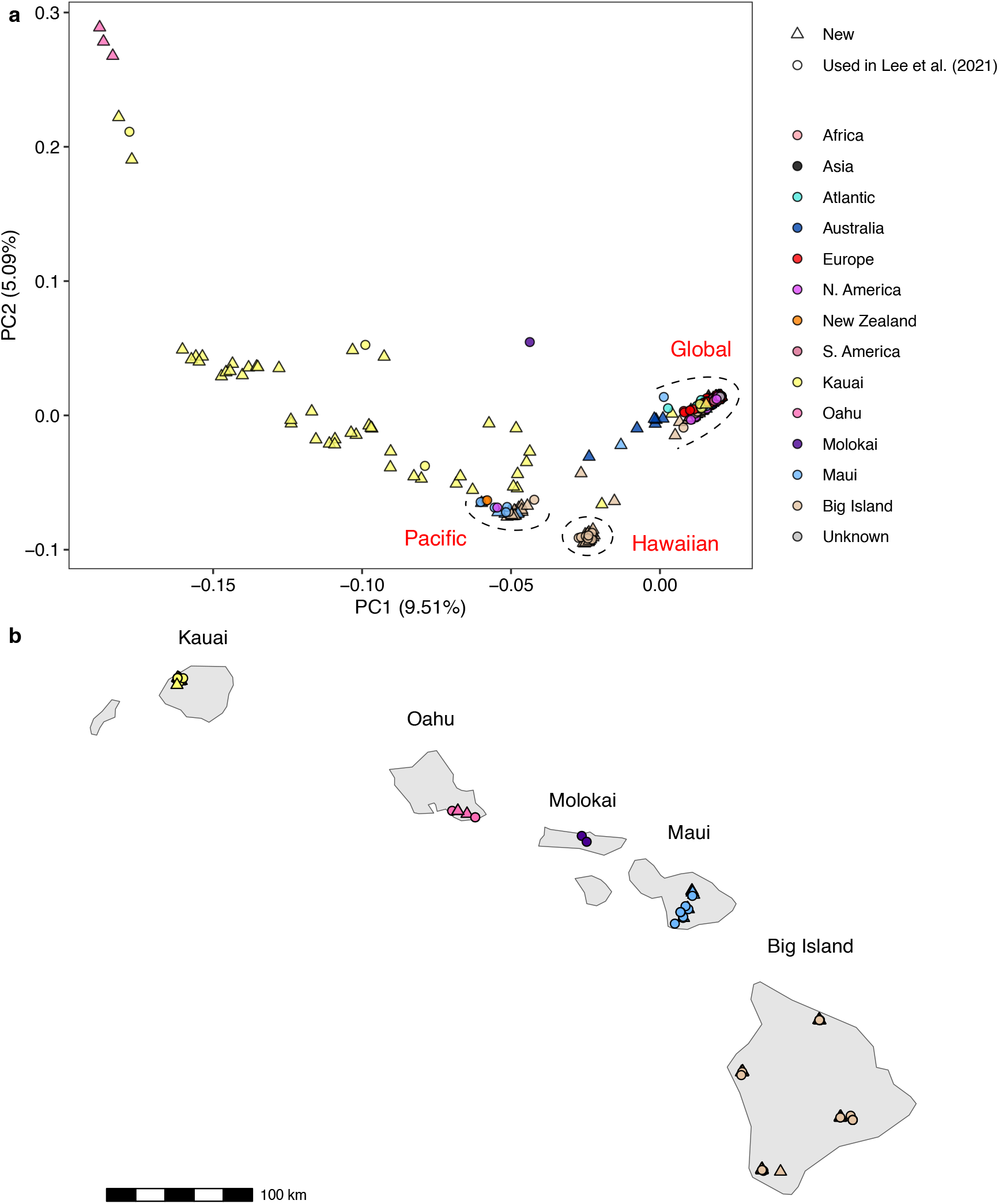
(a) Principal component analysis (PCA) plots of 611 *Caenorhabditis elegans* isotype reference strains based on the genetic variation of variants under the threshold of fraction of missingness (F_MISSING) *<*0.1 with no linkage disequilibrium (LD) pruning, and (b) the geographical locations of Hawaiian strains. Dashed circles or curves delineate three genetic groups that are in line with the ‘global’, ‘Hawaiian’ and ‘Pacific’ groups reported in Lee *et al*. (2021).

When the site-level hard filter is applied, 4,505,918 variants remain (Supplementary Table 1), which is 68.06% of the total variants. The proportion of sites with heterozygous genotypes shrinks to 48.99% in the unpolarized dataset and to 42.03% in the polarized one. Still, most of the sites have fewer than 10 heterozygous genotype calls (Supplementary Fig. 1). There are 3.39 and 2.80 heterozygous calls per site on average after the site-level hard filter for the unpolarized and polarized dataset, respectively (Supplementary Table 1). In summary, despite the prevalence of variants with heterozygous genotypes, most of them are called heterozygous in only a small proportion of strains. Additionally, the site-level hard filter removes heterozygous genotypes more substantially compared to the SNV polarization, indicating that over half of the heterozygous genotypes are of low quality and should be excluded. As a result, we used the hard-filtered dataset with heterozygous sites already removed for the following population structure analysis.

### Population structure analysis

The hard-filtered dataset with all heterozygous genotypes converted to ‘missing’ was used for population structure analyses. In order to retain informative variants while exploring the effect of missing sites and strong linkage disequilibrium (LD) due to the highly selfing nature of the species, we investigated how different filtering thresholds of missing genotype fractions and LD measurements affected the principal component analysis (PCA). At the site level, we further filtered the hard-filtered dataset to remove variants where a proportion of genotypes had missing data within strains. This value is denoted F_MISSING (i.e., fraction of missing genotypes) in BCFtools, and we applied cutoffs from 0 to 0.95 using BCFtools. For the dataset only containing variants with F_MISSING *<*0.1, we applied a series of LD squared correlation (*r*^2^) cutoffs (0.8, 0.6, 0.2 and 0.1) using the `–indep-pairwise` command in PLINK 2.0 (v2.00a5LM) (Chang et al. 2015), with a window size of 50 variants and a step size of 10 variants.

We used VCFtools v.0.1.16 (Danecek et al. 2011) to generate PED/MAP format files from the VCF file as the input of PCA. PCA was then plotted with different F_MISSING and LD pruning cutoffs using the smartpca tool in EIGENSOFT v.8.0.0 (Patterson et al. 2006), which only used biallelic SNVs. The following parameters were used for the PCA without outlier removal: `altnormstyle:NO` ; `numoutevec:50` ; and `familynames:NO` . As the complete-site dataset produces quite different PCA patterns from datasets with missing genotypes (Supplementary Fig. 2), we also ran smartpca with the `missingmode:YES` option for the dataset with the F_MISSING *<*0.1 filter and no LD pruning. This performs PCA on whether each variant has a missing genotype call or not in each sample, which helps to detect structured missingness. We also used VCFtools to calculate the missingness rate of each strain. We calculated the sequencing depth of each strain using mosdepth v.0.2.6 (Pedersen and Quinlan 2017), and the count of missing genotypes that were not called homozygous originally in each strain, indicated by the ‘HP’ field in the VCF file. This step was to determine whether any high levels of missingness were caused by low sequencing depths, or heterozygous SNV polarization that introduced missing genotypes. The correlation between the first principal components under two different PCA modes, and their correlations with F_MISSING were investigated. We used PLINK 2.0 to select strains from the Hawaiian group to run group-specific PCA.

**Figure 2.**
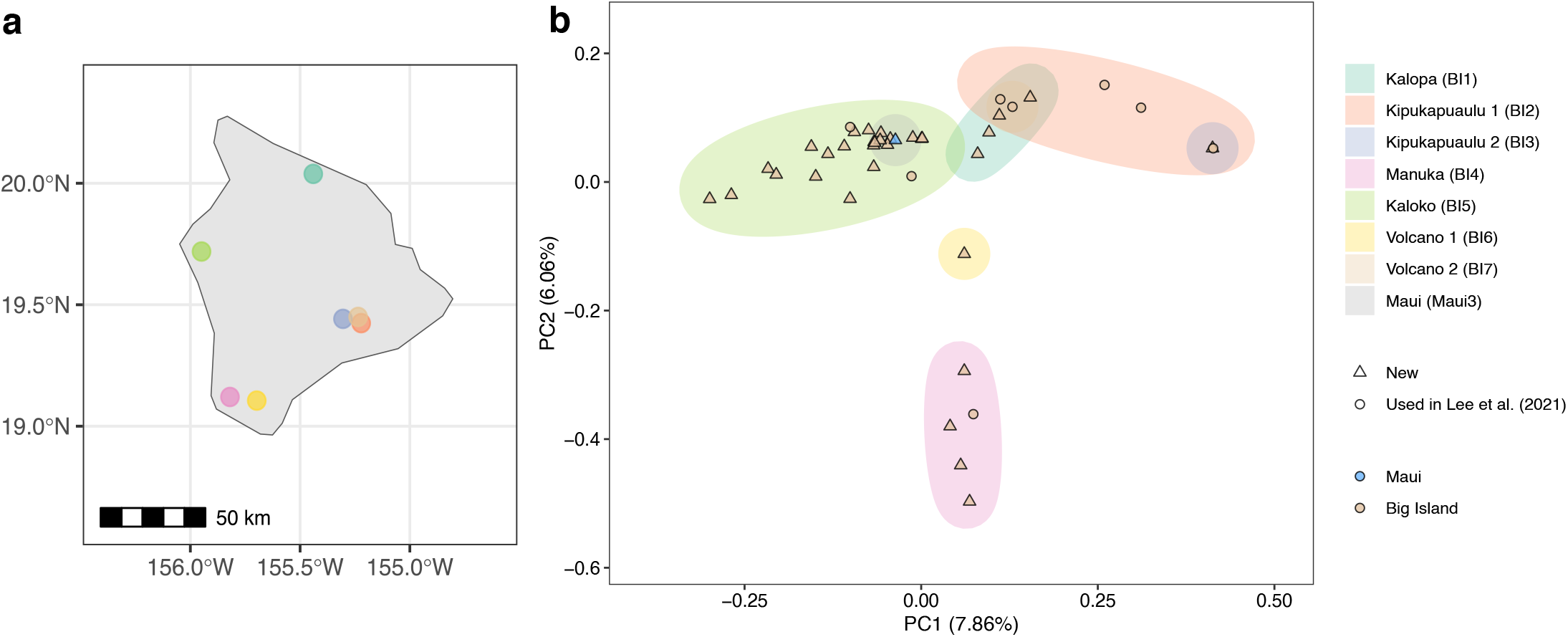
(a) Sampling locations on the Big Island of Hawaii and (b) PCA plot of 41 *C. elegans* strains from the Hawaiian group. Sampling locations defined in Crombie *et al*. (2022) are shown in brackets.

### Demographic inference

There are seven sampling locations with a three-kilometer diameter from the Big Island defined in Crombie *et al*. (2022). Here we named them Big Island Kalopa, Kipukapuaulu 1, Kipukapuaulu 2, Manuka, Kaloko, Volcano and Volcano2, which corresponds to the Big Island 1 to 7 in Crombie *et al*. (2022). We inferred the past demography of strains from four of them–Kalopa, Kipukapuaulu 1, Manuka and Kaloko, which are distributed across the island.

eSMC2 (Sellinger *et al*. 2020) requires multihetsep (MHS) format as input, which records genotypes of segregating sites across samples on each chromosome. We downloaded BAM files of individual strains from CaeNDR to generate MHS files for four subpopulations separately following the MSMC Tools documentation (https://github.com/stschiff/msmc-tools). Specifically, single nucleotide variants (SNVs) were called from BAM files using `bcftools mpileup` and `bcftools call` for each strain. Then bam-Caller.py in MSMC Tools was used to generate single-chromosome VCF files and BED files that encoded the regions meeting coverage threshold for each individual (hereafter coverage masks). These regions were determined based on the mean depth of mapping to each chromosome from each strain calculated using SAMtools v.1.19.2 (Danecek *et al*. 2021). Specifically, regions with depth above twice of the mean and below half of the mean were filtered out. We also generated BED files providing chromosome regions where short sequencing reads can be uniquely mapped. This was done by extracting k-mer subsequences from the reference genome (WS283 release) using `splitfa` from seqbility (Li 2010) and aligning all subsequences back to the genome using BWA v.0.7.18 (Li and Durbin 2009). Then mappability masks were generated using `gen_raw_mask.pl` and `gen_mask` from seqbility and the MSMC Tools script makeMappabilityMask.py.

To investigate how many heterozygous genotypes are likely to be errors, we calculated the number of heterozygous genotypes per 100 kbp window in each strain before and after masking. Specifically, we generated intersections of mappability masks and strain-specific coverage masks for each population using BED-tools v.2.31.1 (Quinlan and Hall 2010). Masks were applied using VCFtools and per-windowed counts were calculated using a customized R script (R Core Team 2023). The distribution of heterozygous genotypes across the genome shows that applying these masks appreciably reduces the number of heterozygous genotypes (Supplementary Fig. 3), suggesting that many of the heterozygous genotypes were artifacts due to mapping errors. This check is compatible with the previous heterozygous site check, which revealed an overall low quality of heterozygous genotypes. Given that result, we excluded variants that were called as heterozygous in these single-strain VCF files. MHS files, which contain the information of segregating sites in regions on which all samples have adequate coverage and that can be uniquely mapped, were generated using `generate_multihetsep.py` from MSMC Tools. Note that no missing genotype data are in the MHS files because they were generated from single-strain VCF files.

**Figure 3.**
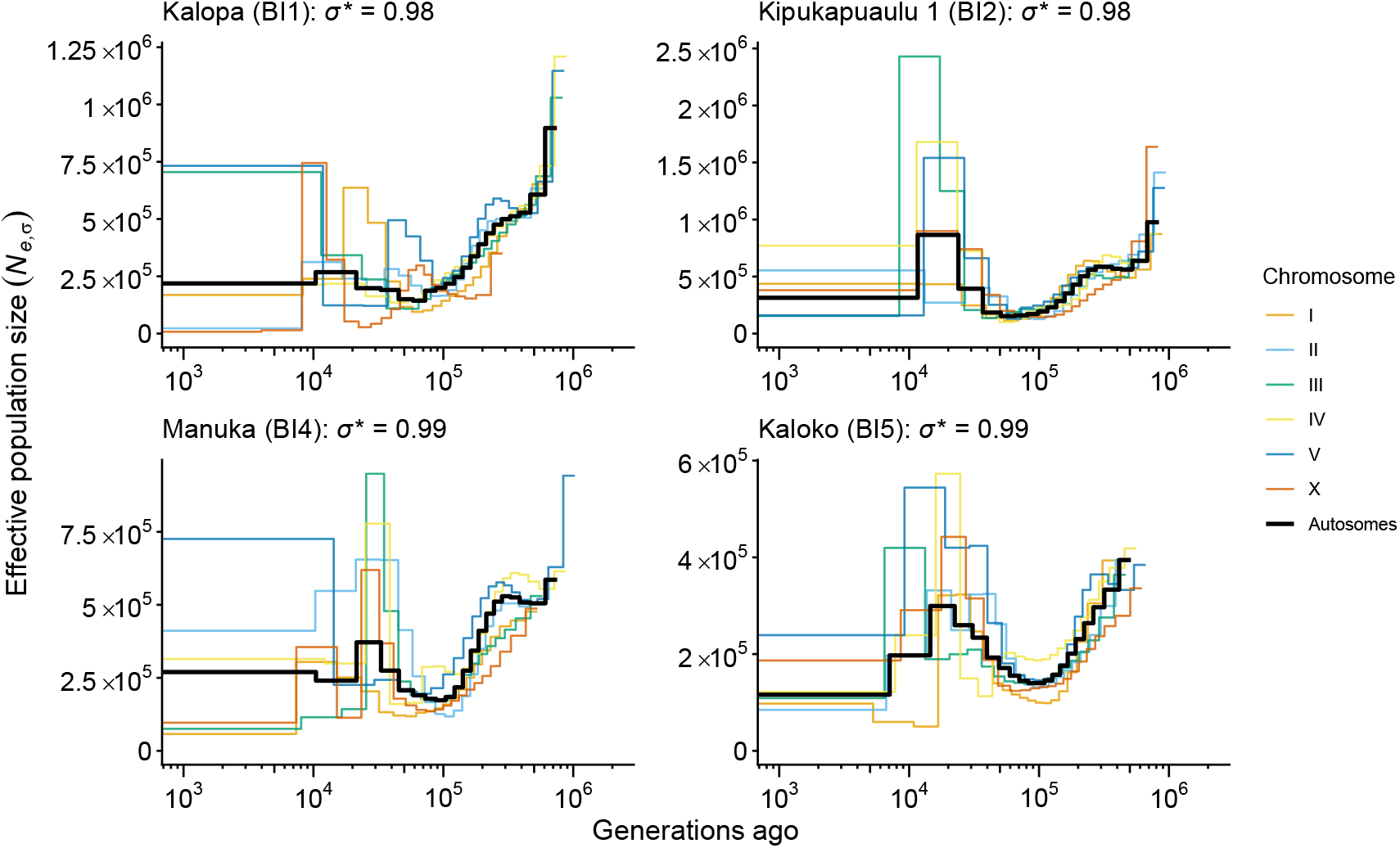
Demographic history and selfing rate for different *C. elegans* populations from Hawaii, as inferred using eSMC2. *σ*^*∗*^ represents the estimated selfing rates. Sampling locations defined in Crombie *et al*. (2022) are shown in brackets. The size estimate more recent than 10^3^ generations is omitted, as no variation of size is inferred later than 10^3^ generations.

We then preserved only one allele copy for each site in each individual strain to generate haploid input for joint inference of demographic history and selfing rate using eSMC2 v.5.3.1 (Sellinger et al. 2020). eSMC2, which assumes a constant selfing rate throughout time, was run on both each chromosome and a combination of all autosomes. We set the number of hidden states (n) to `40` , and BoxP (logarithmic boundaries in base 10 for the population size) to `c(3,3)` , which means the population size can grow up to a thousand times and decrease up to a thousand times. We assumed a mutation rate of *µ* of 2.3 *×* 10^*−*9^ base substitution per basepair per generation (Saxena et al. 2018) as the input mutation rate. Because eSMC2 only takes a single input value of *r* per chromosome, we calculated the weighted average recombination rate of each chromosome based on chromosomal domain-specific estimates in Rockman and Kruglyak (2009), to account for the difference in recombination rate of chromosomal tips, arms, and centers (Table 1). For all autosome data, we used the weighted average across all autosomes, which is 3.08 *×* 10^*−*8^ crossover events per base pair per generation (Table 1). We also ran eSMC2 on a mix of samples from four subpopulations mentioned above (strain ECA1284 from Kalopa; ECA723 from Kipukapuaulu 1; ECA1891 from Manuka; ECA1206 from Kaloko) to detect potential population structure. Note that eSMC2 outputs a vector of population size in unit of effective population sizes corrected for selfing. We thus denotes the eSMC2-inferred population size as *N*_*e,σ*_ to reflect its difference with *N*_*e*_ inferred by selfing-unaware methods.

To investigate the effect on demographic inference when selfing is ignored, we also ran PSMC’, a version of MSMC (Schiffels and Durbin 2014) that uses only two haplotypes, which formed the basis of eSMC2 (Sellinger et al. 2020). The essence of eSMC2 inference is to rescale the input ratio of recombination rate over mutation rate and *N*_*e*_ following the results in Nordborg and Donnelly (1997) and Nordborg (2000). The Nordborg and Donnelly (1997) model points out the coalescent rate of a selfing population is increased by 2/(2 *−σ*), where *σ* is the selfing rate. As *N*_*e*_ is inversely related to the coalescent rate, this leads to a decrease of *N*_*e*_ by (2 *−σ*)/2. As the effective mutation rate (*θ*) is defined as 4*Neµ*, where *µ* is the neutral mutation rate, *θ* is also decreased by (2 *−σ*)/2. However, the effective recombination rate (*ρ*), presented as 4*N*_*e*_*r* where *r* is the molecular recombination rate, is reduced more severely as the high level of homozygosity results in more undetectable recombination events. According to Nordborg (2000), *ρ* is reduced by 1 *− σ* (note that a more exact rescaling for high selfing was derived by Roze (2009), but Sellinger *et al*. (2020) used the result from Nordborg (2000)). Overall, this leads to a rescaling of *ρ*/*θ* by a factor of 2(1 *− σ*)/(2 *−σ*). We hence checked if the size inferred by eSMC2 matches that of PSMC’ after rescaling the latter by 2/(2 *−σ*). We tested this rescaling with pairs of strains from four different Big Island subpopulations used above (ECA1276 and ECA1278 from Kalopa; ECA191 and ECA722 from Kipukapuaulu 1; ECA1261 and ECA1286 from Manuka; ECA1202 and ECA1206 from Kaloko) assuming a *σ* of 0.99, using MSMC v.1.1.0. We used the weighted average recombination rate across all autosomes and the empirical mutation rate to calculate the ratio of recombination rate over mutation rate. The rescaled value of this ratio (*∼* 0.265) was used as the input for the parameter `–rhoOverMu` . The same time window setting as in eSMC2 was applied, which is `20*2` , which means every two of 40 time windows (i.e., inferred time intervals) shares the same value of *N*_*e*_. We also reran eSMC2 on these pairs of strains for comparison.

## Results

### Principal component analysis identifies distinct Hawaiian populations

The original hard-filtered dataset contains 3,536,230 variants, of which 2,835,470 were retained after filtering out those with F_MISSING ≥0.05 (Supplementary Fig. 4). When the F_MISSING threshold is set to 0.1, three genetically distinct groups corresponding to the ‘global’, ‘Hawaiian’ and ‘Pacific’ in Lee *et al*. (2021) are identified (Figure 1). Although less stringent thresholds of F_MISSING preserve slightly more variants, clustering is unaffected: all PCAs of data with non-zero F_MISSING filters recovered a similar population structure as previously found (Lee *et al*. 2021) (Supplementary Fig. 2). Strains from Kauai and Oahu are highly divergent from other strains regardless of the F_MISSING threshold used (Figure 1, Supplementary Fig. 2). By contrast, removing all variants with any missing genotypes generates a complete-site dataset that contains only 1,106,695 variants and clusters all but nine strains from Kauai or Oahu together (Supplementary Fig. 2).

**Figure 4.**
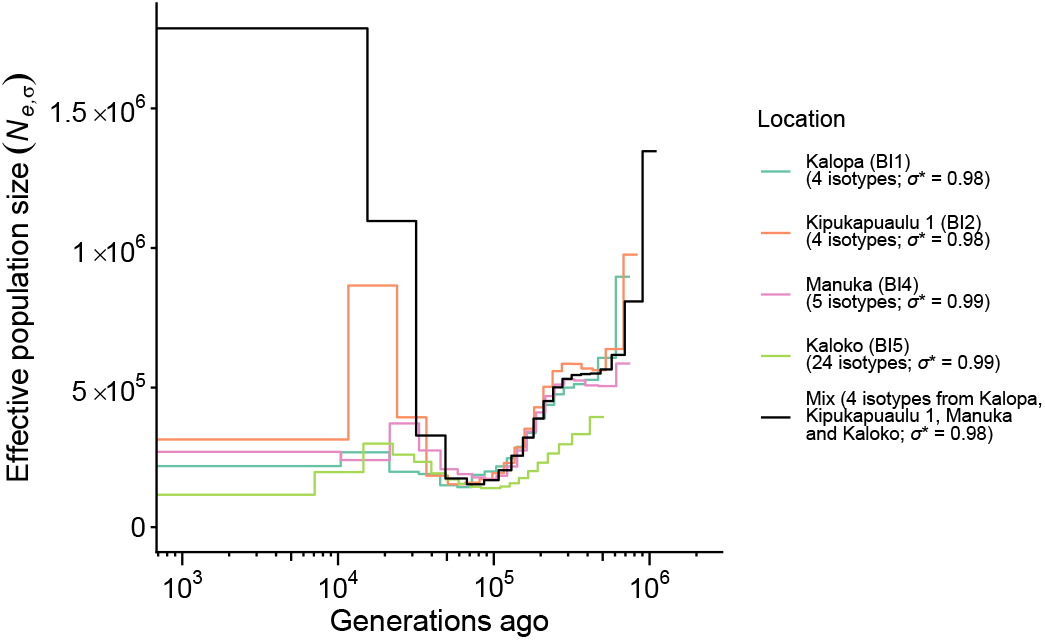
Demographic history and selfing rate estimates in different *C. elegans* populations from Hawaii and a mixed group of strains from different populations, as inferred using eSMC2 and autosome data. Sampling locations defined in Crombie *et al*. (2022) are shown in brackets. The size estimate more recent than 10^3^ generations is omitted, as no variation of size is inferred later than 10^3^ generations.

We thus chose F_MISSING *<*0.1 as the filter before LD pruning, as it preserves population-specific variation patterns (Supplementary Fig. 2).

The PCA based on the variant missingness (with F_MISSING *<*0.1) separates Kauai strains from other strains by the first principal component (PC; Supplementary Fig. 5a). The first PC under this mode is highly correlated with the first PC of the regular PCA mentioned above (Supplementary Fig. 5b) and the first PCs of both PCA are related to a high level of missing genotypes (Supplementary Figs. 5c and 5d). Indeed, we detected more missing genotypes in the Kauai population than other populations, despite the fact that the sequencing depth of Kauai strains is not drastically lower than that of other strains. Although heterozygous SNV polarization can inflate the fraction of missing genotypes, we discover that Kauai strains already have a higher missing rate before heterozygous SNV polarization (Supplementary Fig. 6).

**Figure 5.**
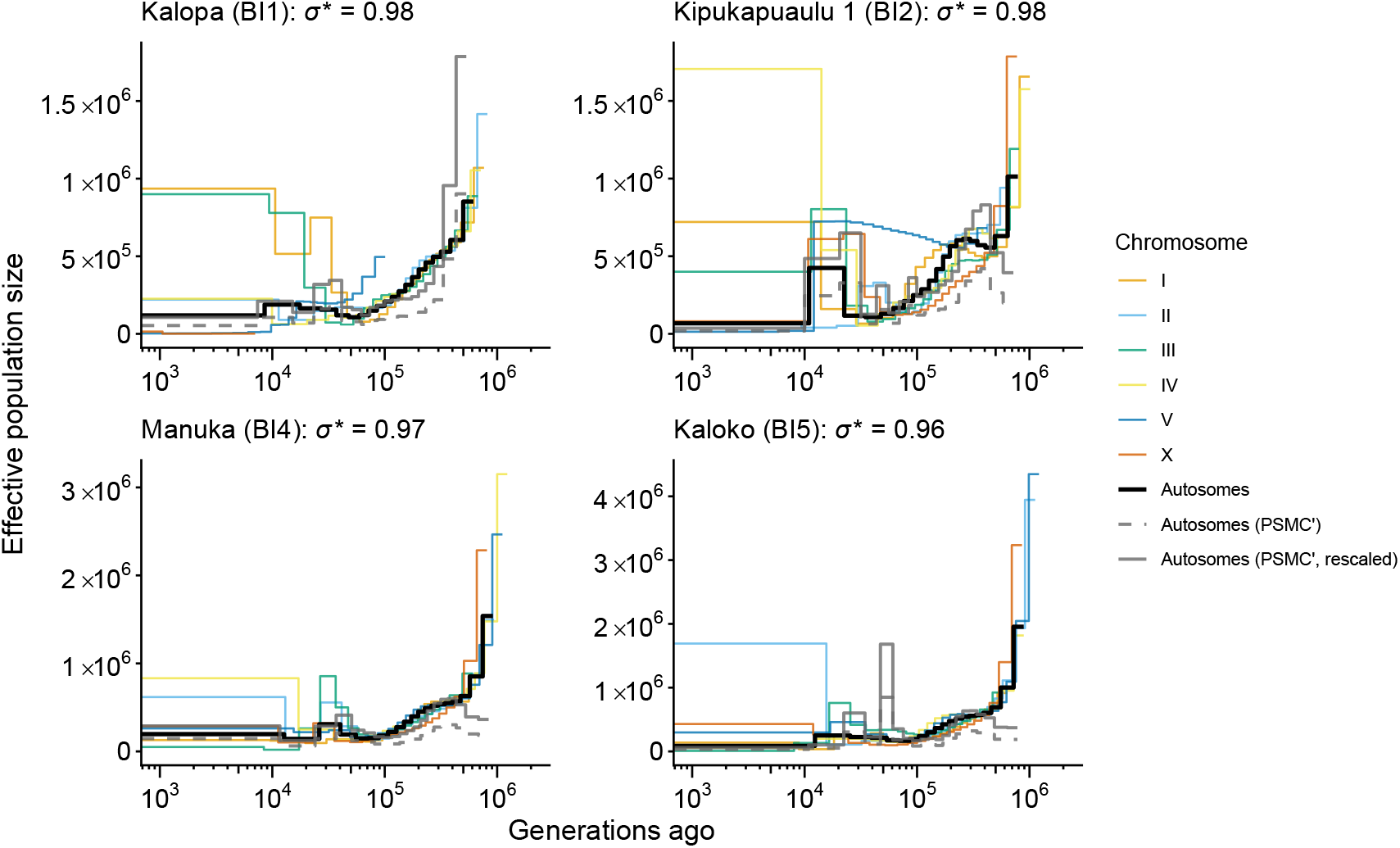
Demographic history in different *Caenorhabditis elegans* populations from Hawaii inferred using eSMC2 and PSMC’. Sampling locations defined in Crombie *et al*. (2022) are shown in brackets. *σ*^*∗*^ represents the selfing rates estimated using eSMC2. The effective population size inferred by eSMC2 accounts for selfing (i.e. *N*_*e,σ*_), while the effective population size originally inferred by PSMC’ doesn’t (i.e. *N*_*e*_). For PSMC’, the original *N*_*e*_ trajectory is represented by dashed grey line and the trajectory rescaled by 2/(2 *−σ*) is represented by solid grey line, where *σ* is the selfing rate (presumed to be 0.99). The size estimate more recent than 10^3^ generations is omitted, as no variation of size is inferred later than 10^3^ generations in most runs (except for the X chromosome run of Kalopa).

LD pruning substantially decreases the number of sites. Even applying a loose LD cutoff (*r*^2^ *<*0.8) removes nearly half of variants, and only about a third of variants remain when a strict cutoff (*r*^2^ *<*0.1) is applied (Supplementary Fig. 7). Removing variants in strong LD largely reduced the genetic divergence among Kauai strains, and between strains from Kauai and the Pacific group (Supplementary Fig. 8). However, the general pattern of three genetic clusters is little affected. Additionally, three Oahu strains and three Kauai strains are genetically distant from the three main clusters, irrespective of the F_MISSING and LD cutoffs used (Supplementary Figs. 2 and 8).

The ‘Hawaiian’ group identified in this study contains 40 strains from the Big Island (collected from 2014 to 2019) and only one from Maui (collected in 2019; Supplementary Table 2). We chose to focus demographic analyses on strains from this group, as they are closely related both genetically and geographically. The group-specific PCA suggests that strains from the same sampling location are clustered together, which implies distinct population structure within the group (Figure 2). We can perform population inference from four of these subpopulations (Kalopa, Kipukapuaulu 1, Manuka and Kaloko) as they contain multiple samples; we hence do so as well as on a dataset that combines samples across locations to investigate the effects of this population structure on inference.

### Demographic and selfing rate inference

eSMC2 estimates selfing rates close to 99% in different Big Island subpopulations from the Hawaiian genetic group using all autosomes (Figure 3). This result is consistent with previous estimates of selfing rate in most *C. elegans* wild populations (Barrière and Félix 2005a,b; Cutter 2006; Barrière and Félix 2007). Selfing rate estimates based on different chromosomes vary little, with all estimates over 96% (Supplementary Table 3). Decreases in *N*_*e,σ*_ from *∼*1 *×* 10^5^ to *∼* 5 *×* 10^5^ generations ago are inferred across all four Big Island subpopulations (Figure 3 and 4). Although demographic trajectories are in line with those previously inferred by Teterina *et al*. (2023), the *N*_*e,σ*_ inferred by eSMC2 are larger by an order of magnitude. *N*_*e,σ*_ changes inferred from different chromosomes are largely identical between 10^5^ and 10^6^ generations ago, with greater variation observed in more recent generations (Figure 3). When we applied the same analysis to four samples from different Big Island subpopulations, we observe drastic *N*_*e,σ*_ increases in the recent past (Figure 4). This result is likely a signal of strong population structure inflating coalescent times, and hence estimates of *N*_*e,σ*_ (Slatkin 1991; Mazet *et al*. 2016). The inferred selfing rate for this mixed sample is about 0.98, with little variation among chromosomes (Supplementary Table 3).

We next tested if accounting for selfing significantly changes the inferred demographic history by comparing the demographic trajectory inferred by eSMC2 and the *N*_*e*_ trajectory inferred by PSMC’ (Figure 5). eSMC2 inferences with only pairs of haplotypes produce similar demographic trajectories as those using all samples. Selfing rate estimates are slightly smaller in general, except for one estimate of 30% for the X chromosome data of Kalopa samples (Supplementary Table 3). This might be a misinference, arising due to too few segregating sites identified when the sample size is decreased (Supplementary Table 3), which substantially changes the inferred demographic history (Figure 5; the inferred size is constantly smaller than 10^5^). Demographic trajectories inferred by PSMC’ also show similar shapes to their eSMC2 counterparts, indicating a population decline. However, its original *N*_*e*_ estimates are smaller than *N*_*e,σ*_ inferred by eSMC2 in general. After rescaling PSMC’-inferred *N*_*e*_ by 2/(2*− σ*) *≈* 1.98 for *σ* = 0.99, the demographic trajectories inferred using PSMC’ are largely consistent with their eSMC2 counterparts.

## Discussion

Inferring the historical population size using genomic data informs us of how populations responded to previous environmental changes and provides a genetically neutral model for detecting selection signals. However, the genetic effect of reproductive mode, in particular self-fertilization, has been largely ignored in popular demographic inference methods, with a few exceptions (Sellinger et al. 2020; Strütt et al. 2023; Peede et al. 2025). In this study, we inferred the demographic history of the Big Island population of *C. elegans* from Hawaii, a predominantly selfing nematode and compared SMC-based methods that either do or do not account for the effect of selfing. Despite its global distribution, the Hawaiian population of the species exhibits a distinct genetic background from global populations (Crombie et al. 2019; Lee et al. 2021). The 2023 release of the CaeNDR dataset includes more strains from different Hawaiian islands, especially the Big Island and Kauai, which allows us to recover a more comprehensive picture of within-island population structure, highlighting the robustness of genetic groups reported in the previous study (Lee *et al*. 2021), and the genetic distinctness of the Kauai population. We find that eSMC2 gives accurate estimates of selfing rate for *C. elegans* and infers a historic population decline that is shared different Big Island subpopulations. By default, *N*_*e,σ*_ estimated by eSMC2 is larger than *N*_*e*_ estimated by PSMC’. However, by rescaling both the ratio of recombination and mutation rate, and the output *N*_*e*_ in PSMC’ to account for self–fertilization, PSMC’ produces similar *N*_*e*_ estimates to eSMC2.

### Distinct genetic diversity pattern of Hawaiian populations

Lee *et al*. (2021) reported three genetically distinct groups of *C. elegans* strains at the global scale: the ‘Hawaiian’ group, which contains strains only from the Big Island of Hawaii (corresponding to the ‘Volcano’ group in Crombie *et al*. (2019)); the ‘Pacific’ group, which mostly consists of strains from Hawaii, but also includes one strain from North America and one from New Zealand (corresponding to the ‘HI Divergent’ group in Crombie *et al*. (2019) excluding a few very divergent strains (ECA701, XZ1514 and XZ1516) from Kauai); and the ‘global’ group, which includes all other strains. Our PCA recovers a Hawaiian group with the majority of strains from the Big Island (Figures 1 and 2, Supplementary Fig. 8). The Pacific group contains strains from both the Big Island and Maui, and the aforementioned two strains outside Hawaii (Figure 1, Supplementary Fig. 8). Strains from Kauai, the oldest sampled Hawaiian island, exhibit more genetic divergence between strains than those from the three main genetic groups when variants with strong LD are not removed (Supplementary Fig. 2). This reflects the higher fraction of missing genotypes in Kauai strains rather than any difference in sequencing quality (Supplementary Figs. 5a and 6). After LD pruning, most of the Kauai strains cluster closely with the Pacific group (Supplementary Fig. 8). This might be attributable to group-specific hyper-divergent haplotypes, as reported for other *C. elegans* populations worldwide (Lee *et al*. 2021). Hyper-divergent regions are defined as regions that have a particularly high variant density or low coverage by Lee *et al*. (2021), corresponding to a decreased mapping rate of short sequence reads to the reference and an increase in missing genotype calls. Future long-read sequencing of Kauai strains is required to investigate whether specific hyper-divergent regions or structural variants accounts for the elevated missing rate.

Additionally, three strains from Kauai (ECA2191, ECA2195 and XZ1516) and three strains from Oahu (ECA1493, ECA1713 and ECA3088) are consistently divergent from other strains regardless of missingness filters and LD pruning thresholds used (Supplementary Fig. 2 and 8). XZ1516 and ECA701 were reported as the most divergent isotypes in Lee *et al*. (2021). One of the two Molokai isotypes (ECA369) also separates from all three genetic groups when there is no LD pruning (Supplementary Fig. 2). These findings suggest that Kauai, Oahu, and Molokai populations could harbor a large amount of distinct variation. More samples from these Hawaiian islands are needed to disentangle when and how *C. elegans* colonised the different islands of the Hawaiian archipelago and to identify the potential underlying drivers.

The PCA of strains from the Hawaiian group shows clustering by sampling locations (Figure 2), suggesting spatial population structure at the scale of *∼* 50 to *∼* 100 km. Strong local population structure is expected in selfing populations as a consequence of both dispersal limitation and extreme inbreeding (Charlesworth *et al*. 1997) and has been reported in non-Hawaiian populations before (Barrière and Félix 2005a; Sivasundar and Hey 2005; Barrière and Félix 2007; Dey 2007). The only Maui strain of the group clusters with strains from the BI5 subpopulation, suggesting that it might be recently introduced from the Big Island (Figure 2).

### Demographic history of the Big Island population

eSMC2 infers a selfing rate of close to 99% with either single chromosome or all autosomes in all four Big Island subpopulations when over four samples are used, which matches estimates in previous studies (Figure 3). When only two samples are used, most estimates are still over 94%. The only exception is inferred from the X chromosome data of Kalopa samples, which contains a very low number of segregating sites (Supplementary Table 3). In terms of demographic inference, even when over four samples are used, *N*_*e,σ*_ inferred from the X chromosome are still different from those inferred from autosomes in Kalopa (Figure 3), which may again result from fewer usable segregating sites (Supplementary Table 3). We conclude that, for extreme selfers, it is prefereable to use data from multiple chromosomes and/or samples to avoid noisy estimates if there are a low number of segregating sites. Different autosomes generate similar demographic trajectories in general, though greater variation is seen when inferring recent *N*_*e,σ*_ . This variation is expected given the limited resolution in the recent past when considering pairwise coalescence (Li and Durbin 2011; Sellinger et al. 2020). Population declines from about *∼*1*×* 10^5^ to about *∼*5*×* 10^5^ generations ago were inferred in different subpopulations (Figures 3 and 4). Previous demographic analyses also report a similar decline when running PSMC on pairs of strains outside Hawaii (Thomas et al. 2015), or when running SMC++ on a Big Island population with a ‘global’ genetic background (Teterina et al. 2023). It is likely that the ‘Hawaiian’ and ‘global’ groups share the same ancestral population during the decline, which requires further investigation of the split time between two groups. *N*_*e,σ*_ inferred here is also larger than *N*_*e*_ inferred in Thomas *et al*. (2015) and Teterina *et al*. (2023). This might be in part be due to the fact that Thomas *et al*. (2015) ignored the effects of selfing (while Teterina *et al*. (2023) rescale both mutation rate and coalescent time), but could also reflect using only strains from the ‘global’ population, which has lower genetic diversity (Crombie et al. 2019), in their analyses. Assuming a generation time of 25 effective generations per year (Wei et al. 2026), our analyses suggest a population decline from 20 thousand years ago (kya) to 4 kya. However, the driver of this population decline remains elusive, as does the geographic location of the ancestral population.

By mixing strains from different sampling locations on the Big Island, we detected an inferred increase of *N*_*e,σ*_ since about 15 thousand generations ago (Figure 4). Although the *N*_*e,σ*_ of the Kipukapuaulu 1 subpopulation peaked around the same time, none of the separated subpopulation runs reached a plateau of *N*_*e,σ*_ increase as the mixed subpopulation run. Instead, we conclude that the large recent *N*_*e,σ*_ inferred for the mixed sample is an artifact of strong population structure (Li and Durbin 2011; Mazet *et al*. 2016). It highlights the importance of defining subpopulations for demographic inferences, as sampling genomes from different locations may lead to false signatures of *N*_*e,σ*_ (or *N*_*e*_) change, which is a common misinterpretation in SMC-based demographic inference (Bansal and Nichols 2025; Hilgers *et al*. 2025). This step is especially important for self-fertilising species which are more likely to be sub-structured (Charlesworth *et al*. 1997).

### The importance of accounting for selfing in demographic inferences

Comparing the results between eSMC2 and PSMC’ illustrates the consequence of ignoring the effect of selfing in demographic inferences. Overall smaller sizes are inferred using PSMC’ in all Big Island subpopulations as its *N*_*e*_ estimates assume a randomly mating, outcrossing poulation. However, perhaps unsurprisingly the inferred trajectory of *N*_*e*_ change follows the same shape as *N*_*e,σ*_ inferred using eSMC2 and the difference between the two analyses disappears when we rescale the PSMC’-inferred *N*_*e*_ by 2/(2 *− σ*). Note that although rescaling these parameters manually when using PSMC’ can generate similar results, this can only be done when the selfing rate is known. eSMC2 is therefore a better choice when there is limited information about the selfing rate of the population.

Despite being able to correct the inferred histories of population size change in selfing populations, we stress that biases are still expected between estimates of *N*_*e,σ*_ and census population size. Both eSMC2 and PSMC’ do not account for the effect of linked selection. Using synonymous substitutions in genic regions or masking regions potentially under selection are common strategies for eliminating the impact of selection (Marchi et al. 2021). However, in selfing populations the effect of linked selection is amplified due to lower effective recombination rates (Golding and Strobeck 1980; Nordborg 2000; Hartfield and Glémin 2014, 2016; Hartfield et al. 2017; Hartfield and Glémin 2024). While Hawaiian populations of *C. elegans* do not exhibit the global selective sweep that underpins the dominant haplotype observed in non-Hawaiian populations (Andersen et al. 2012; Lee et al. 2021), background selection is still likely to reduce genetic diversity (and hence *N*_*e*_) further (Charlesworth et al. 1993; Barrett et al. 2014). The effect of background selection in reducing *N*_*e*_ can be stronger when the selfing rate is over 90% (Glémin and Ronfort 2013; Barrett et al. 2014). Recent simulation-based studies have also demonstrated how strong linked selection may distort the shape of demographic trajectories inferred by SMC-based methods. For example, Johri *et al*. (2021) shows that a population decline can be falsely inferred as growth when the proportion of directly selected sites in the genome increases to 20%. Crucially and unlike demographic history, linked selection results in different rates of genetic drift affecting different segments of the genome, creating heterogeneity in *N*_*e*_ across the genome (Charlesworth 2009; Gossmann et al. 2011; Jiménez-Mena et al. 2016). Thus inferred demographic trajectories can be strongly biased by extensive linked selection and, in particular, lead to a decrease in inferred *N*_*e*_ even when the population sizes have remained constant (Boitard et al. 2022). Developing methods that account for the effect of linked selection is still essential for improving the accuracy of demographic inferences for selfing populations, and will facilitate us to uncover the historical demographic changes experienced in *C. elegans* both on and outside Hawaii.

## Conclusion

Self-fertilization can lead to some profound genetic effects in demographic inference, which may cause a misinterpretation of the inferred demographic trajectory and is often ignored in SMC-based inferences. Here we used eSMC2, an SMC-based method to jointly infer the selfing rate and the historical population size changes of the Big Island population of *C. elegans*, a predominantly self-fertilizing nematode, to investigate how it accounts for the effect of selfing in demographic inference. We recovered reliable selfing rate estimates and a gradual population decline reported in previous studies on other *C. elegans* populations. We obtained population size estimates that are in general twice of the *N*_*e*_ inferred using PSMC’, a selfing-unaware method, and show that this difference can be reconciled using a simple rescaling of *N*_*e*_ estimates under the assumption of a constant selfing rate. Our analyses also suggests that Big Island–specific strains show similar population declines to populations outside Hawaii, and that clear population structure exists among different Big Island subpopulations. Our work paves the way for building more realistic background models for *C. elegans* which not only capture its demographic history but also account for the joint effects of selfing, past demography and linked selection on patterns of genetic diversity.

## Data availability

The code used in this work is available at https://github.com/Wangeos/Celegans_demography.

