## Supplementary Materials for "Selfing-aware demographic inference highlights local population structure and bottlenecks in Hawaiian *Caenorhabditis elegans*"

2026-09-22

### Supplementary Tables

Table 1: Heterozygous genotype-related statistics for *Caenorhabditis elegans* datasets with different filters.

Table 2: Sample information of isotype reference strains from the 'Hawaiian' group.

Table 3: Individual counts, number of segregating sites and the estimated selfing rate ($\sigma^*$) of each eSMC2 run.

### Supplementary Figures

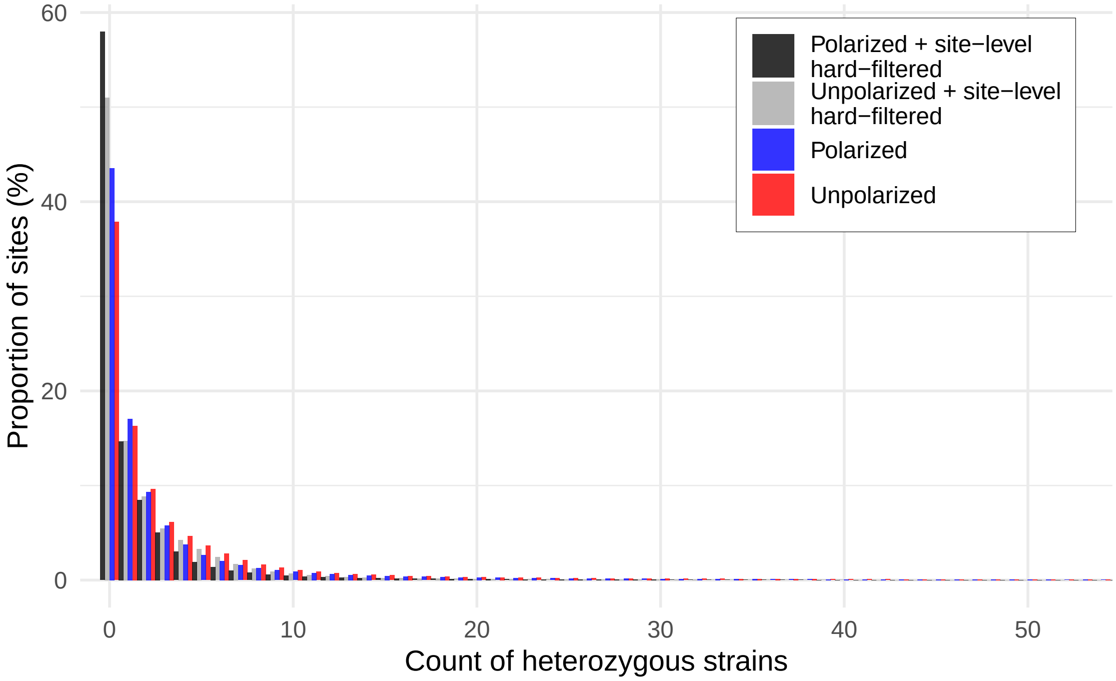


Figure 1: The distribution of heterozygous genotype count of all variants in the unpolarized, polarized, unpolarized and site-level hard-filtered, polarized and site-level hard-filtered datasets of *Caenorhabditis elegans*. Entries with more than 50 heterozygous strains are omitted as sums of them are smaller than 5% in all four datasets.

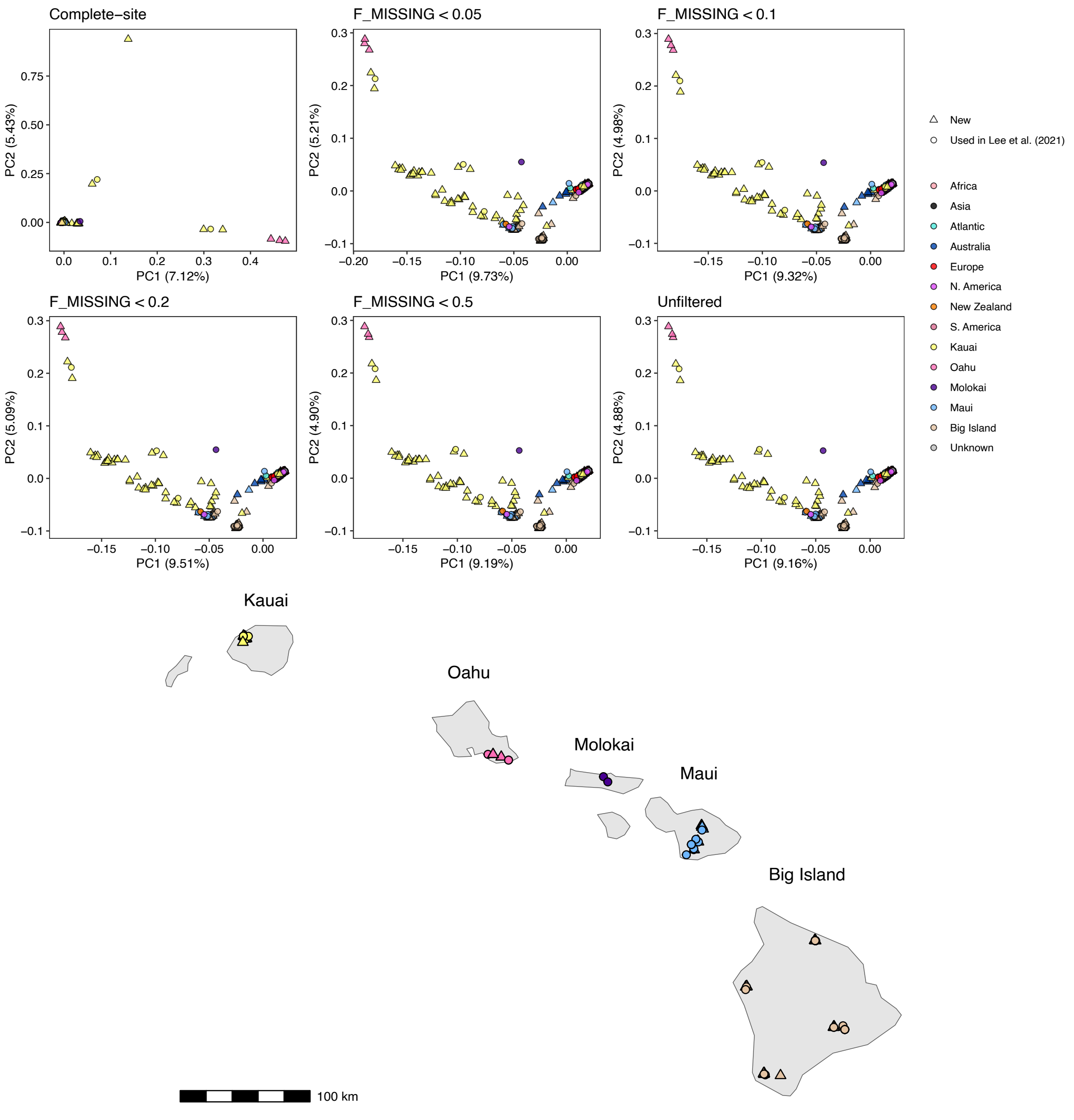


Figure 2: Principal component analysis (PCA) plots of 611 *Caenorhabditis elegans* strains based on the genetic variation of variants under different fraction of missingness (F_MISSING) thresholds, and the geographical locations of Hawaiian strains. 'Complete-site' refers to the threshold where variants that are missing in any strains are filtered out.

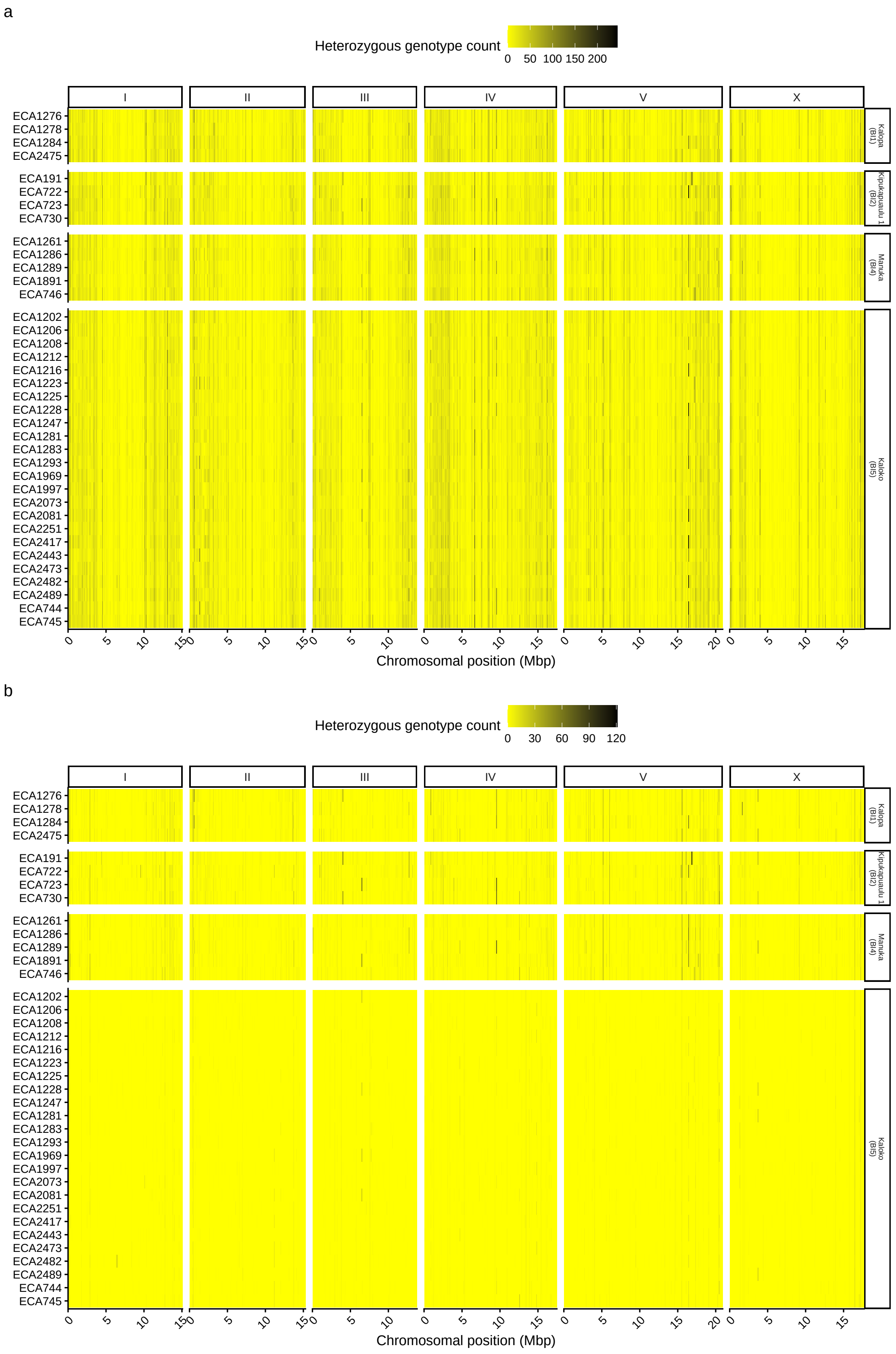


Figure 3: The heterozygous genotype counts per 100-kbp window with a step of 10 kbp, calculated from single-strain VCF files of all strains used in eSMC2 (a) before and (b) after coverage and mappability masking. Sampling locations defined in Crombie *et al.* (2022) are shown in brackets.

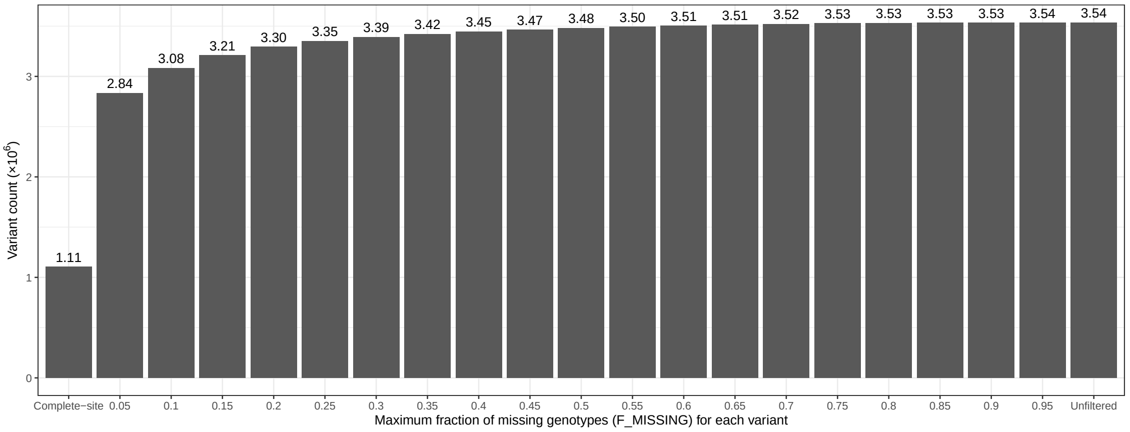


Figure 4: Number of variants passing the fraction of missing genotype (F_MISSING) filter from the hard-filtered dataset. 'Complete-site' refers to the threshold where variants that are missing in any strains are filtered out.

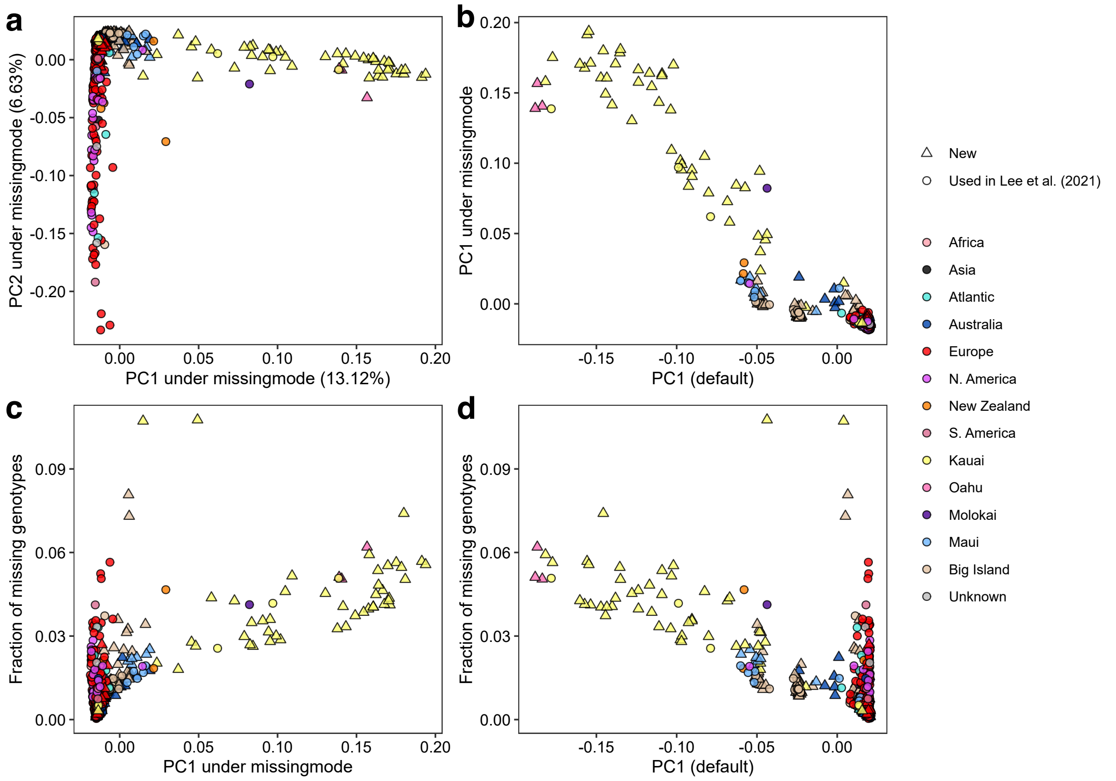


Figure 5: (a) PCA plots of 611 *Caenorhabditis elegans* strains based on whether each variant has a missing genotype call or not in each sample. The variant dataset with the fraction of missing genotypes (F_MISSING) $< 0.1$ filter and no LD pruning is used. (b) Correlation between the first principal components (PC1) of default PCA and PCA under missingmode. (c) Correlation between the PC1 of PCA under missingmode and fraction of missing genotypes in each strain. (d) Correlation between the PC1 of default PCA and fraction of missing genotypes in each strain.

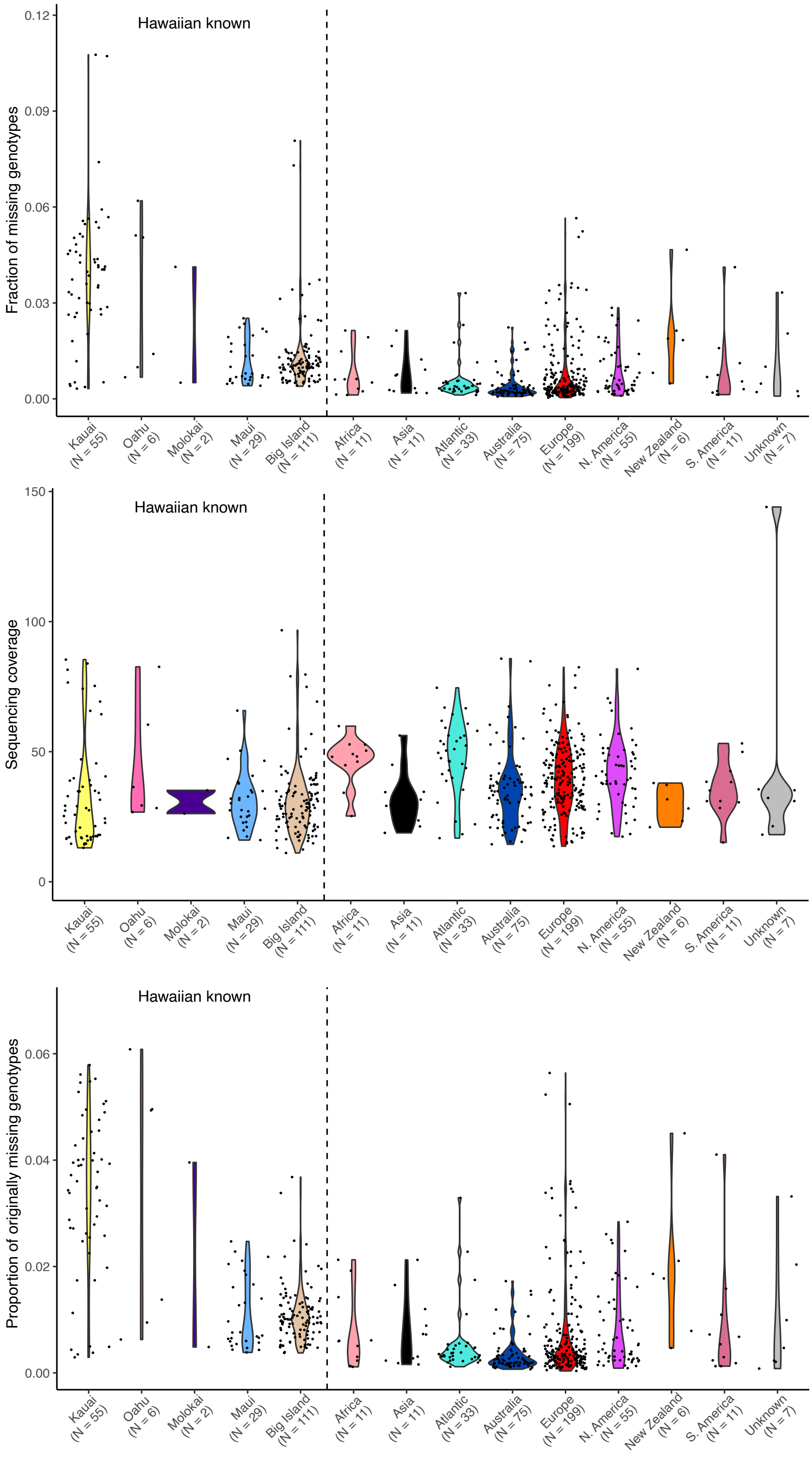


Figure 6: The proportion of missing genotypes in each strain, sequencing coverage and the proportion of originally missing calls of strains before heterozygous SNV polarization from different geographical regions. The variant dataset with the fraction of F_MISSING $< 0.1$ filter and no LD pruning is used. N, sample size.

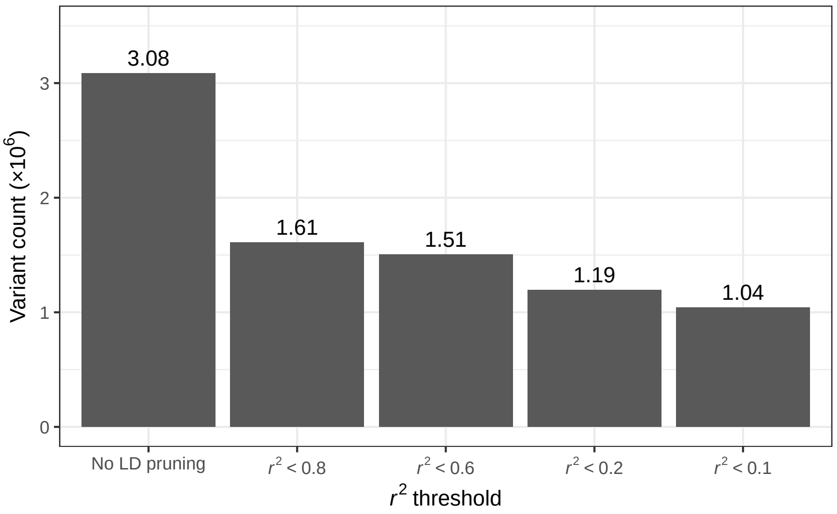


Figure 7: Number of variants passing the $r^2$ filter from the dataset of sites with F_MISSING < 0.1.

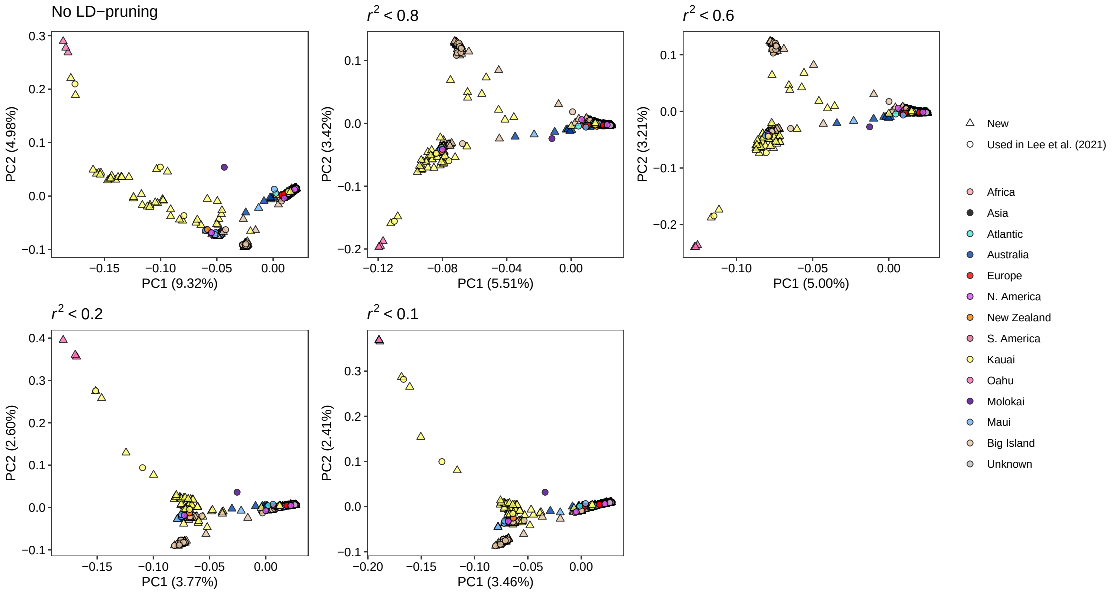


Figure 8: Principal component analysis (PCA) plots of 611 *Caenorhabditis elegans* strains based on the genetic variation of variants under different LD pruning thresholds. LD pruning was applied to the dataset with F_MISSING < 0.1.
